# White Matter Hyperintensity Quantification in Large-Scale Clinical Acute Ischemic Stroke Cohorts – The MRI-GENIE Study

**DOI:** 10.1101/552844

**Authors:** Markus D. Schirmer, Adrian V. Dalca, Ramesh Sridharan, Anne-Katrin Giese, Kathleen L. Donahue, Marco J. Nardin, Steven J. T. Mocking, Elissa C. McIntosh, Petrea Frid, Johan Wasselius, John W. Cole, Lukas Holmegaard, Christina Jern, Jordi Jimenez-Conde, Robin Lemmens, Arne G. Lindgren, James F. Meschia, Jaume Roquer, Tatjana Rundek, Ralph L. Sacco, Reinhold Schmidt, Pankaj Sharma, Agnieszka Slowik, Vincent Thijs, Daniel Woo, Achala Vagal, Huichun Xu, Steven J. Kittner, Patrick F. McArdle, Braxton D. Mitchell, Jonathan Rosand, Bradford B. Worrall, Ona Wu, Polina Golland, Natalia S. Rost, on behalf of the MRI-GENIE Investigators

## Abstract

White matter hyperintensity (WMH) burden is a critically important cerebrovascular phenotype linked to prediction of diagnosis and prognosis of diseases, such as acute ischemic stroke (AIS). However, current approaches to its quantification on clinical MRI often rely on time intensive manual delineation of the disease on T2 fluid attenuated inverse recovery (FLAIR), which hinders high-throughput analyses such as genetic discovery.

In this work, we present a fully automated pipeline for quantification of WMH in clinical large-scale studies of AIS. The pipeline incorporates automated brain extraction, intensity normalization and WMH segmentation using spatial priors. We first propose a brain extraction algorithm based on a fully convolutional deep learning architecture, specifically designed for clinical FLAIR images. We demonstrate that our method for brain extraction outperforms two commonly used and publicly available methods on clinical quality images in a set of 144 subject scans across 12 acquisition centers, based on dice coefficient (median 0.95; inter-quartile range 0.94-0.95) and Pearson correlation of total brain volume (r=0.90). Subsequently, we apply it to the large-scale clinical multi-site MRI-GENIE study (N=2783) and identify a decrease in total brain volume of −2.4cc/year. Additionally, we show that the resulting total brain volumes can successfully be used for quality control of image preprocessing.

Finally, we obtain WMH volumes by building on an existing automatic WMH segmentation algorithm that delineates and distinguishes between different cerebrovascular pathologies. The learning method mimics expert knowledge of the spatial distribution of the WMH burden using a convolutional auto-encoder. This enables successful computation of WMH volumes of 2,533 clinical AIS patients. We utilize these results to demonstrate the increase of WMH burden with age (0.950 cc/year) and show that single site estimates can be biased by the number of subjects recruited.

## Introduction

White matter hyperintensity (WMH) burden is a clinically important and highly heritable cerebrovascular phenotype (Atwood et al., 2004; Debette and Markus, 2010). Utilizing magnetic resonance imaging (MRI), WMH can be readily identified on T2 fluid attenuated inverse recovery (FLAIR) images due to the increased contrast (DeCarli et al., 2005). FLAIR is a common MRI sequence used for clinical assessment in acute ischemic stroke (AIS) patients. Additionally, WMH burden has been linked to cerebrovascular disease outcomes (Wardlaw et al., 2013), especially in ischemic stroke (Smith, 2010), where the underlying genetic effects are still largely unknown (Giese et al., 2017). Manual or semi-automatic methods for delineating WMH are labor intensive and time consuming, making them impractical in large-scale studies. Fully automatic approaches are therefore necessary to enable genetic discovery based on image-derived phenotypes in large-scale studies.

Recently, several groups have presented automatic algorithms to identify and differentiate WMH from other hyperintense signals on T2 MRI (Caligiuri et al., 2015; Dadar et al., 2017). Most algorithms are developed using research quality scans with isotropic, or close to isotropic, resolution. Additionally, these methods require significant (often multimodal) preprocessing of the images, such as brain extraction and spatial normalization to a template (Caligiuri et al., 2015), which is particularly challenging with clinically acquired data. When imaging patients hospitalized with acute stroke, isotropic sampling is infeasible as the increased acquisition time would interfere with demands of acute clinical care. The resulting images have high in-plane, but low through-plane resolution, due to large slice thickness and/or spacing between slices. Subsequently, image analysis steps essential for automatic WMH delineation often fail, and therefore dedicated workflows (also called computational pipelines) are required to accommodate clinical images (Schirmer et al., 2017; Sridharan et al., 2014, 2013).

Segmenting WMH in AIS populations is particularly challenging (Dalca et al., 2014), as the T2 hyperintense stroke lesion may also be visible on the FLAIR sequence, and are not separated from the underlying WMH burden by most algorithms. Therefore, manual or semi-automatic protocols have been used to segment WMH in AIS patients (Cloonan et al., 2015; Etherton et al., 2017). These approaches preclude large-scale analysis.

In this work, we develop a high-throughput, fully automated WMH analysis pipeline for clinical grade FLAIR images, which promises to facilitate rapid phenotypic evaluation and demonstrate our pipeline in a large-scale multi-site study of AIS patients. The pipeline incorporates: (a) brain-extraction specifically designed for clinical FLAIR images, (b) intensity normalization to accommodate for multi-site heterogeneity, and (c) automatic atlas-based segmentation of WMH in the image. We introduce a new algorithm for brain extraction, utilize a mean-shift algorithm for intensity normalization, and build on previously demonstrated methods for WMH segmentation in stroke patients (Dalca et al., 2014). We validate the efficacy of this pipeline on a carefully curated validation image set and demonstrate the pipeline efficacy in a large multi-site study of AIS patients, where we analyze WMH volume (WMHv) associations with age in a cross-sectional AIS study cohort.

## Methods

### Neuroimaging data

The MRI-GENetics Interface Exploration (MRI-GENIE) study is a large-scale, international, hospital-based collaborative study of AIS patients (Giese et al., 2017). Approval from the institutional review board or ethics committee was obtained by each site in line with their institutional guidelines. Informed consent was obtained for all patients by the individual sites. FLAIR scans of 2,781 patients from 12 sites (7 European, 5 US based) were acquired over a period of more than eight years, ending 2011, as part of each hospital’s clinical AIS protocol. The axial FLAIR images have a mean resolution of 0.7mm in-plane (minimum: 0.4mm, maximum: 1.9mm) and 6.3mm through-plane (minimum: 1.0mm, maximum: 65.0mm). Additionally, basic demographics, such as age and sex, are available for all subjects in the study. Table A. summarizes the MRI-GENIE data set used in our experiments.

### Validation set

For each acquisition site, we selected 12 subjects approximately spanning the range of WMH volume based on qualitative assessment, forming a validation image set (N=144, Table A.1). For each subject in the validation set, the brain and WMH were segmented manually. ‘Manual’ outlines were performed based on a semi-automated method with high inter-rater reliability (Chen et al., 2006; Gurol et al., 2006), after undergoing a structured training protocol for WMH segmentation, which demands a high inter-rater agreement (Intra-class Correlation Coefficient (ICC) of 0.92). Here, a single rater (K.D.) who passed the training protocol and with over 2 years of experience has completed the WMH segmentation. We use these segmentations for quantitative evaluation of automatic analysis steps. Table A. also summarizes the validation set. Methods are developed without access to the validation set, which is only used once the development concluded in order to demonstrate/validate the efficacy of the developed methods in previously unseen data.

### Pipeline Overview

Each image undergoes brain extraction and intensity normalization, followed by the WMH segmentation. The WMH segmentation algorithm identifies white matter disease using atlas-based prior models for spatial distribution of WMH and local intensities in the image (Dalca et al., 2014). We perform the necessary spatial normalization to an age-appropriate FLAIR template via the affine registration implementation in the ANTs software package (Avants et al., 2011). Figure 1 presents an overview of the fully automatic analysis pipeline for WMH segmentation in clinical FLAIR images. Implementation was based on a large-scale processing infrastructure to enable processing of thousands of scans within parallel deployment systems (Sridharan et al., 2013).

**Figure 1:**
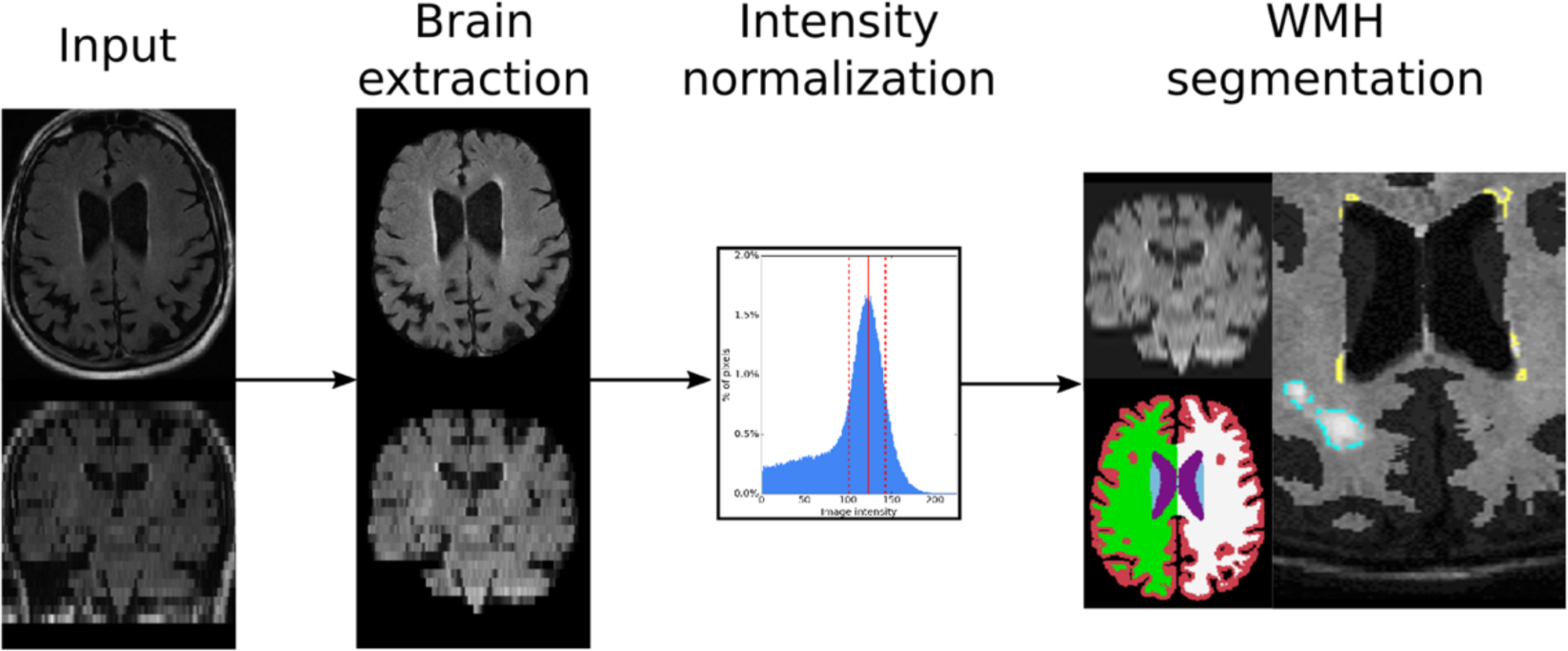
Overview of the analysis pipeline for extracting WMH in clinically acquired FLAIR images. Each input image first undergoes brain extraction, followed by intensity normalization. Images are spatially normalized, i.e. upsampled and affinely registered to an atlas, in order to allow for WMH segmentation with spatial priors.

### Brain extraction

We develop a brain extraction method for clinical FLAIR scans (Neuron-BE) that employs a 2D U-Net convolutional neural network architecture^1^ (see schematic in Figure 2). We first roughly normalize the intensities of each image, so that the 97^th^ percentile of the image intensities is scaled to 1 and pad the image so that the in-plane resolution is a multiple of 16. The architecture contains five downsampling levels and five upsampling levels. Down-/up-sampling is achieved using 2×2 maxpool/upsample operations. Each level contains two convolution layers with 128 features per layer. To optimize the network parameters, we use the Adadelta stochastic optimizer (Zeiler, 2012) with mini-batches of size 16. For each batch, we augment the data to mimic the observed conditions in clinical data. The augmentation includes random intensity scaling (contrast factor between 0.7 – 1.3), random ghosting effects (at most 3 “copies” of the brain), as well as additive Gaussian and Perlin noise (standard deviations of 0.4 and 0.5, respectively). We learn the network parameters using a supervised training set of 69 subject scans for which we manually outlined the brain. These manual outlines include grey matter, white matter, and the four ventricles, and were generated by tracing the outer boundary between grey matter and CSF. Given learned parameters, we apply our convolutional neural network to a subject’s FLAIR image. Additionally, we close holes in the resulting segmentation mask, and identify the largest connected component as the brain mask (Van der Walt et al., 2014).

**Figure 2:**
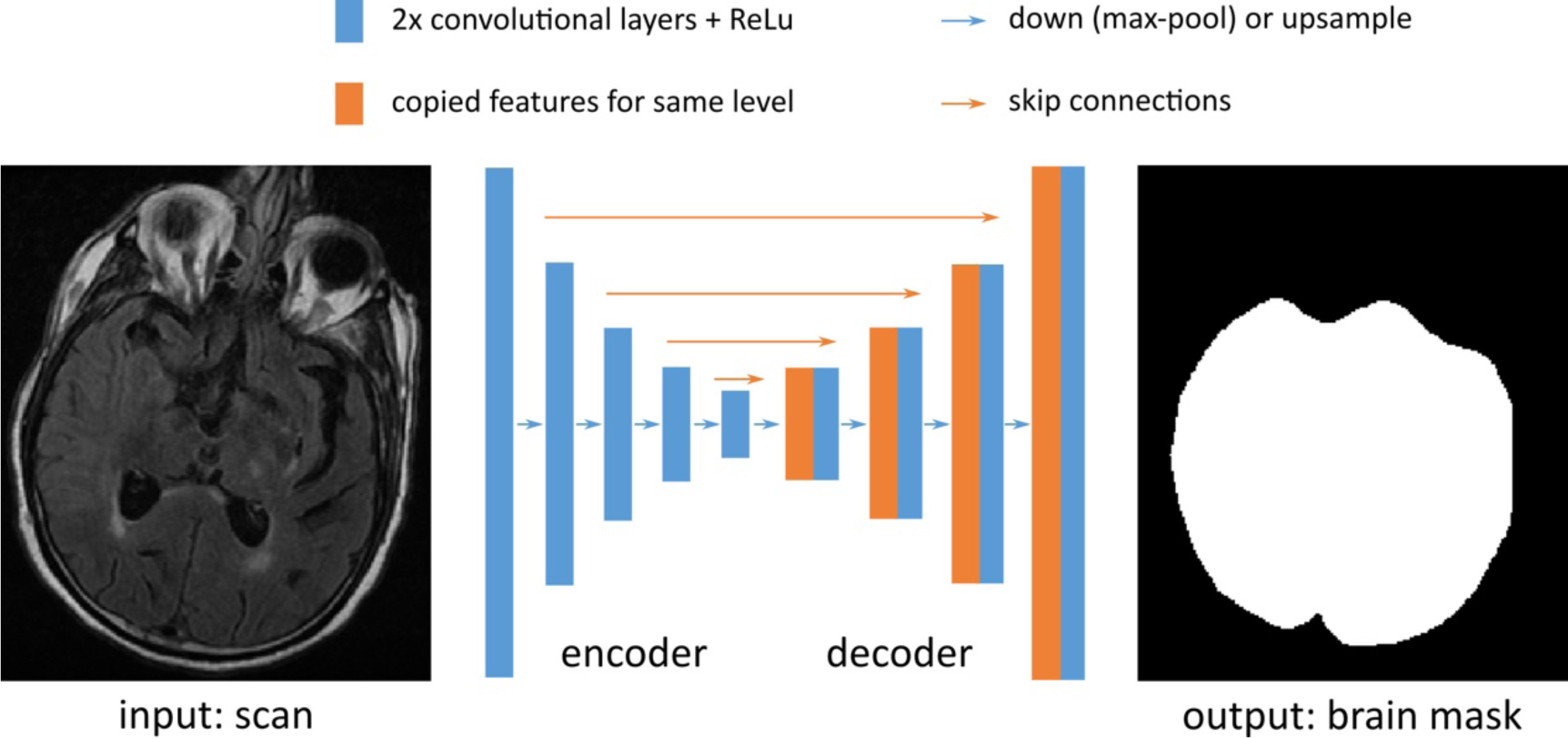
The Neuron-BE architecture, based on the UNet, contains five downsampling levels and five upsampling levels, achieved using 2×2 maxpool/upsample operations (blue arrows). Each level contains two convolution layers with 128 features per layer. To optimize the network (convolution) parameters, we use the Adadelta stochastic optimizer with mini-batches of size 16.

To assess the efficacy of the brain extraction on clinical scans, we compare results of Neuron-BE and two publicly available methods (ROBEX (Iglesias et al., 2011) and FSL BET (Smith, 2002)), within our validation set. For each resulting segmentation, we compute volume overlap (via Dice coefficient (Dice, 1945)) and correlation of total volumes estimates (via Person’s correlation coefficient and after outlier removal based on the modified z-score on the volume differences) between manually and automatically generated brain masks.

### Intensity normalization

Tissue intensity values vary substantially across FLAIR scans. Intensity normalization is therefore useful for harmonization across scanners and imaging sites. However, WMH can lead to failure of traditional histogram normalization (Sridharan et al., 2013). Therefore, after brain extraction we use a mean-shift algorithm (Cheng, 1995) to determine the mode of intensity distribution that corresponds to the average white matter intensity in each scan (Sridharan et al., 2013). Brain extraction is an essential first step for intensity normalization, to ensure that “background” intensities, originating e.g. from the skull, eyes or neck, are not taken into account in the estimation of the mode. Image intensity values are rescaled, so that the mode maps to an intensity of 0.75.

We assess intensity normalization using the validation set by using the intensity value estimated via the mean-shift algorithm and the corresponding full width half maximum (FWHM) of the peak in the intensity histogram of the total brain volume to mark potential white matter voxels. These outlines can be used to confirm visually that the majority of voxels in these masks correspond to white matter, enabling qualitative visual assessment of the intensity normalization in each subject of the validation set. Additionally, we visually assess the cumulative white matter masks of the 144 subjects in template space after affine registration.

### Automatic WMH segmentation

We build on the an existing automatic WMH segmentation algorithm (Dalca et al., 2014) to delineate and distinguish different cerebrovascular pathologies on brain MRI. The algorithm is derived from a generative probabilistic model that describes T2 FLAIR image intensities of WMH and stroke lesions. The model captures disease priors using a 2D convolutional auto-encoder that mimics experts’ knowledge of the spatial distribution of WMH (see schematic in Figure 3). The auto-encoder contains four sets of convolution layers, max-pooling and down-sampling layers, a dense layer to capture spatial covariance and create a fixed-length encoding, and four sets of convolution and up-sampling layers. It uses the ReLu activation function on all convolution layers. We optimize the parameters of the neural network using an independent set of manual WMH outlines (Zhang et al., 2015; 699/91/90 outlines used for training/validating/testing) via stochastic updates with the Adadelta optimizer (Zeiler, 2012).

**Figure 3:**
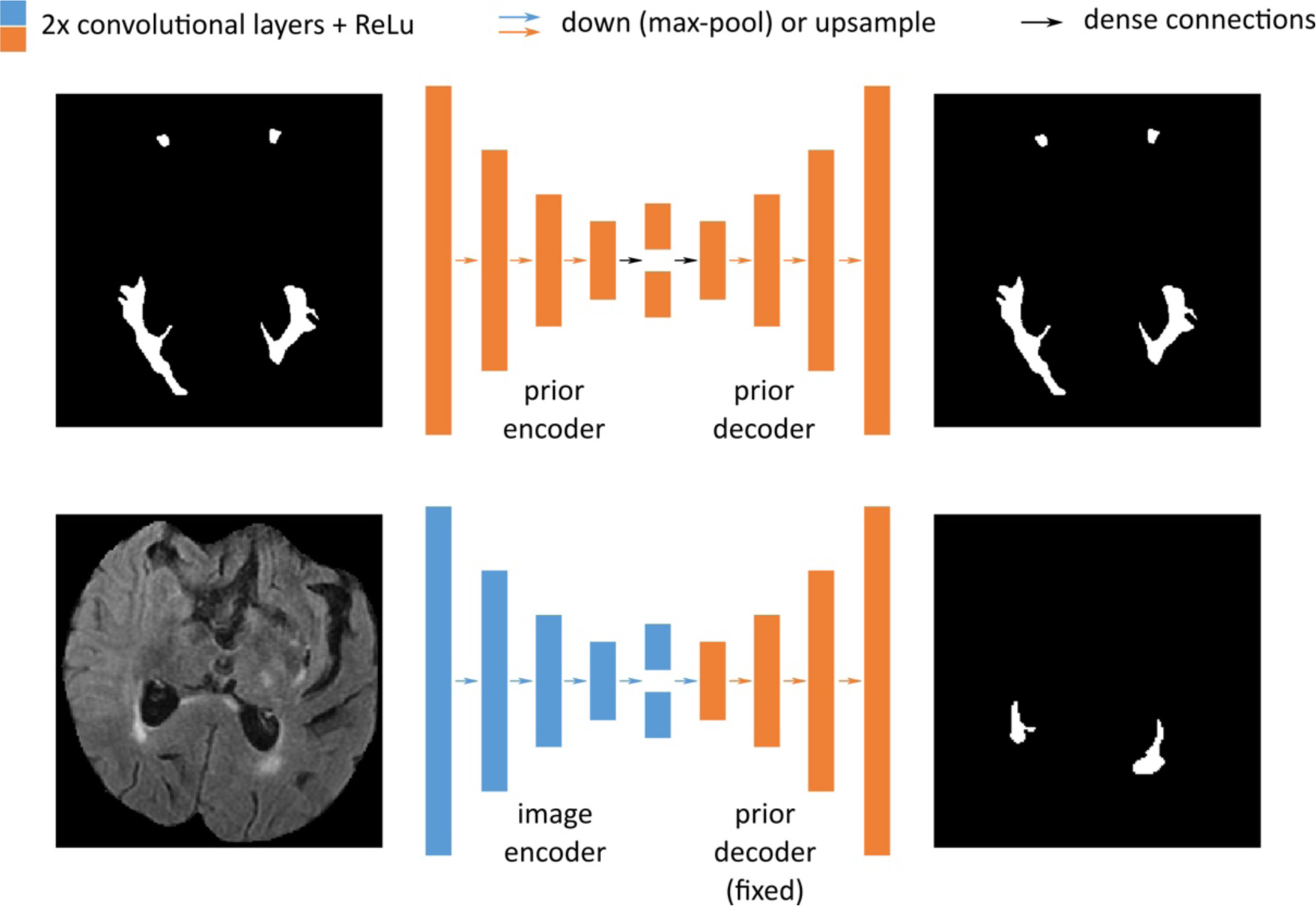
Architecture for automated WMH segmentation. The model first captures disease priors using a convolutional auto-encoder (top) that mimics experts’ knowledge of the spatial distribution of WMH. The auto-encoder contains four sets of convolution layers, max-pooling layers, a dense layer (black arrow) to capture spatial covariance and create a fixed-length encoding, and four sets of convolution and up-sampling layers. We use ReLu activation function on all convolution layers. The prediction network (bottom) uses this (fixed) prior by taking an input scan and projecting down to an encoding using a similar architecture as above with independent parameters, before using the prior decoder weights to yield a segmentation from this encoding.

In order to employ spatial priors, we interpolate each input scan using bi-cubic upsampling (Jones et al., 2014) and register the upsampled image to age-appropriate FLAIR template (1mm^3^, dimensions: 182, 218, 182 in x, y, and z; Schirmer et al., 2018) using affine alignment (Avants et al., 2011). We assess registration efficacy by calculating the Dice coefficient between each subject’s brain mask and the brain mask of the template.

Additionally, we evaluate the efficacy of the automated WMHv quantification on the validation data set using Pearson’s correlation coefficient and the ICC as measures of agreement with manual segmentations. Furthermore, we assess the residuals (variations of the automatically estimated WMHv from the manual) for trends based on the age and sex phenotypes.

### Quality control (QC) in the MRI-GENIE image set

As a first step of QC, we determine outliers based on both the in-plane and through-plane resolution for all scans using the modified z-score with a threshold of 3.5 (Iglewicz and Hoaglin, 1993). Additionally, we flag scans with low slice numbers (≤3). The detected potential outliers are flagged for visual evaluation, where the user can determine if they represent valid data or if they should be excluded from subsequent analyses.

After brain extraction, we detect and eliminate potential outliers by examining the modified z-scores computed on total brain volume based on the extracted brain masks. Here, total brain volume is defined as grey matter, white matter, and the four ventricles. In addition to the site-based modified z-score (threshold 3.5), we model brain volume changes with age as

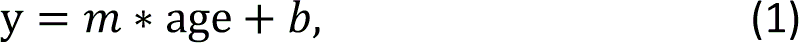

where *y* is the brain volume, and *m* and b the regression parameters. Parameters are estimated using the Python package *numpy (Ascher et al., 1999)*. We use this model and the standard deviation of its residuals, to determine age-based outliers. Outliers are defined as brain volumes more than two standard deviations away from the model estimate. Similar to the initial QC step with respect to the image resolution, we flag subjects for manual QC, which helps detect potential erroneous WMH segmentations due to incomplete brain extraction. Finally, after processing the remaining images, three raters randomly select two WMH outlines per site and qualitatively assess the quality of the resulting segmentations.

### WMHv analysis in MRI-GENIE image set

We investigate the extracted WMHv for each individual site and compare them to the pooled data set, comprising all subjects of all sites, by computing the distributions of WMHv. In addition, we estimate the coefficient of change in WMHv with age for the pooled data set and investigate the effect of sample size. To do so, we model the association of the natural log-transformed WMHv as a linear function of age (see equation (1)). Additionally, we calculate uncertainty (standard deviation) of the determined coefficients of change for each site using a 10-fold split of the data for each site and a subsequent leave-one-fold-out approach.

## Results

### Validation set

We first characterize the distribution of volumes of WMH obtained from manual delineation in the validation set. Figure 4 (left) shows the histogram of WMHv across subjects (0.24-119.98 cc).

**Figure 4:**
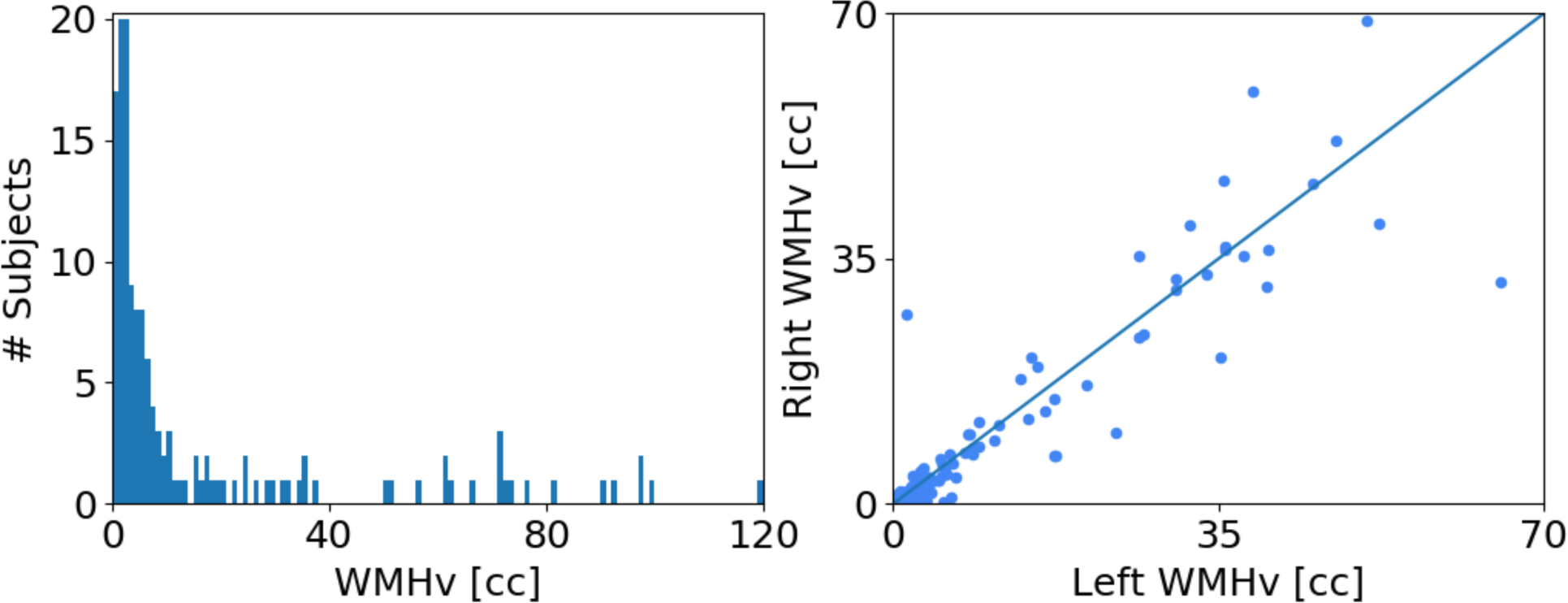
WMHv in the validation set based on manual segmentations of WMH (144 subjects, 12 per site). Left: Distribution of WMHv. Right: Comparison of left and right hemispheric WMHv (Wilcoxon: p<0.05).

### Brain extraction

Figure 5 shows the volume overlap between the automatically extracted brain mask and the manual brain segmentation for ROBEX (Iglesias et al., 2011), BET (Smith, 2002) and Neuron-BE on the validation set scans.

**Figure 5:**
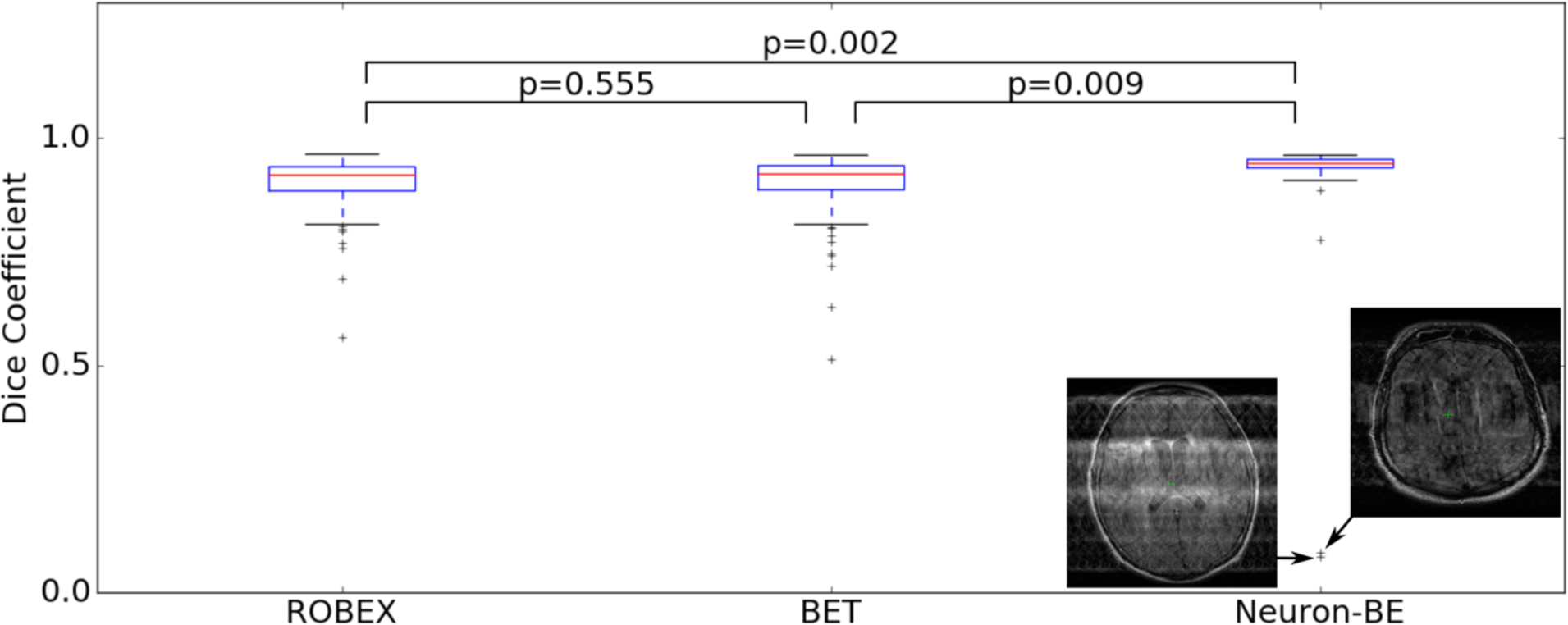
Volume overlap distributions between the automatically extracted brain mask and a manual brain segmentation in the validation set for ROBEX, FSL BET and Neuron-BE. Comparisons between methods are based on paired t-tests. Median Dice coefficient were 0.92, 0.92 and 0.95 for ROBEX, FSL BET and Neuron-BE, respectively.

Median volume overlap (inter-quartile range (IQR)), as measured by the Dice coefficient, was 0.92 (0.94-0.89), 0.92 (0.94-0.89) and 0.95 (0.95-0.94) for ROBEX, BET and Neuron-BE, respectively. We remove outliers using the modified z-score on the volume difference between total volume estimates obtained from automatic segmentations and those extracted from manual segmentations (3, 8 and 14 subjects for Neuron-BE, ROBEX and FSL BET). Correlations were subsequently estimated to be 0.94, 0.80 and 0.75 for Neuron-BE, ROBEX and FSL BET (p<0.01 for all correlations), respectively.

Neuron-BE also helps us identify outliers with substantial imaging artefacts. We find two gross outliers in the brain extraction using Neuron-BE due to motion and ghosting effects. Severe motion corruption or ghosting effects can prevent accurate WMH segmentation. We therefore flag these scans as potentially problematic for WMH segmentation. Computing z-scores for outlier detection identifies both of these scans as outliers in the case of NEURON-BE, but not for ROBEX or FSL BET. Upon visual inspection, the remainder of the scans did not show any motion corruption or ghosting artefacts.

Overall, our results demonstrate that Neuron-BE offers a more appropriate approach to brain extraction than popular publicly available methods. For the remainder of the analysis we use Neuron-BE and incorporate a quality control step based on brain volume. This enables us to obtain accurate brain extraction results on clinical data while being able to remove images where resulting WMH segmentations are ill-defined due to image quality.

### Intensity normalization

Figure 6 (a) shows an example of the estimated white matter intensity using the mean shift algorithm, and full-width-half-maximum (FWHM) of the distribution peak. The resulting “white matter mask” is shown in Figure 4 (b). Finally, Figure 4 (c) shows the cumulative white matter mask in atlas space.

**Figure 6:**
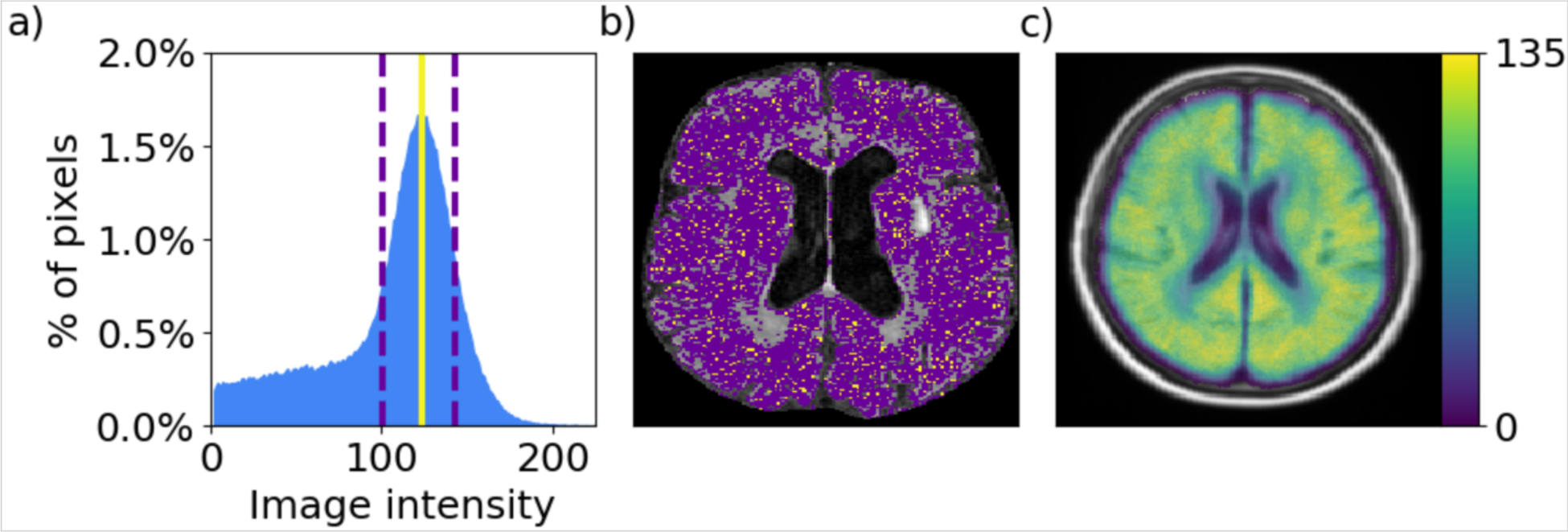
a: Example intensity distribution for one subject with the estimated mean white matter intensity (solid yellow line) and full width half maximum (FWHM; dashed lines). b: Axial slice of the corresponding FLAIR image. Voxels whose image intensity is equal to the estimated mean white matter intensity are shown in yellow; voxels whose image intensity falls into FWHM range are shown in purple. c: Cumulative white matter mask in atlas space for all 144 subjects.

Visual inspection of the white matter masks on all 144 subjects of the validation set and the cumulative white matter mask suggest that the average white matter intensity estimates are accurate and can be used to normalize image intensity across sites. We normalize each subject’s FLAIR image intensities, by scaling the intensity distribution so that the mean white matter intensity equals 0.75.

### Automatic WMH segmentation

We spatial normalize the clinical scans by first upsampling each scan, and affinely registering the result image to the FLAIR atlas. The Dice coefficient of the brain masks of each affinely registered subject scan compared to the brain mask of the template were 0.93±0.01 (mean ± standard deviation). We apply the automatic WMH segmentation algorithm to each affinely registered scan. Figure 7 shows the comparsion of WMHv (ICC=0.84, Pearson r = 0.86 with p<0.001) between manual and automatic outlines (left), as well as the residuals (right), for the validation set.

**Figure 7:**
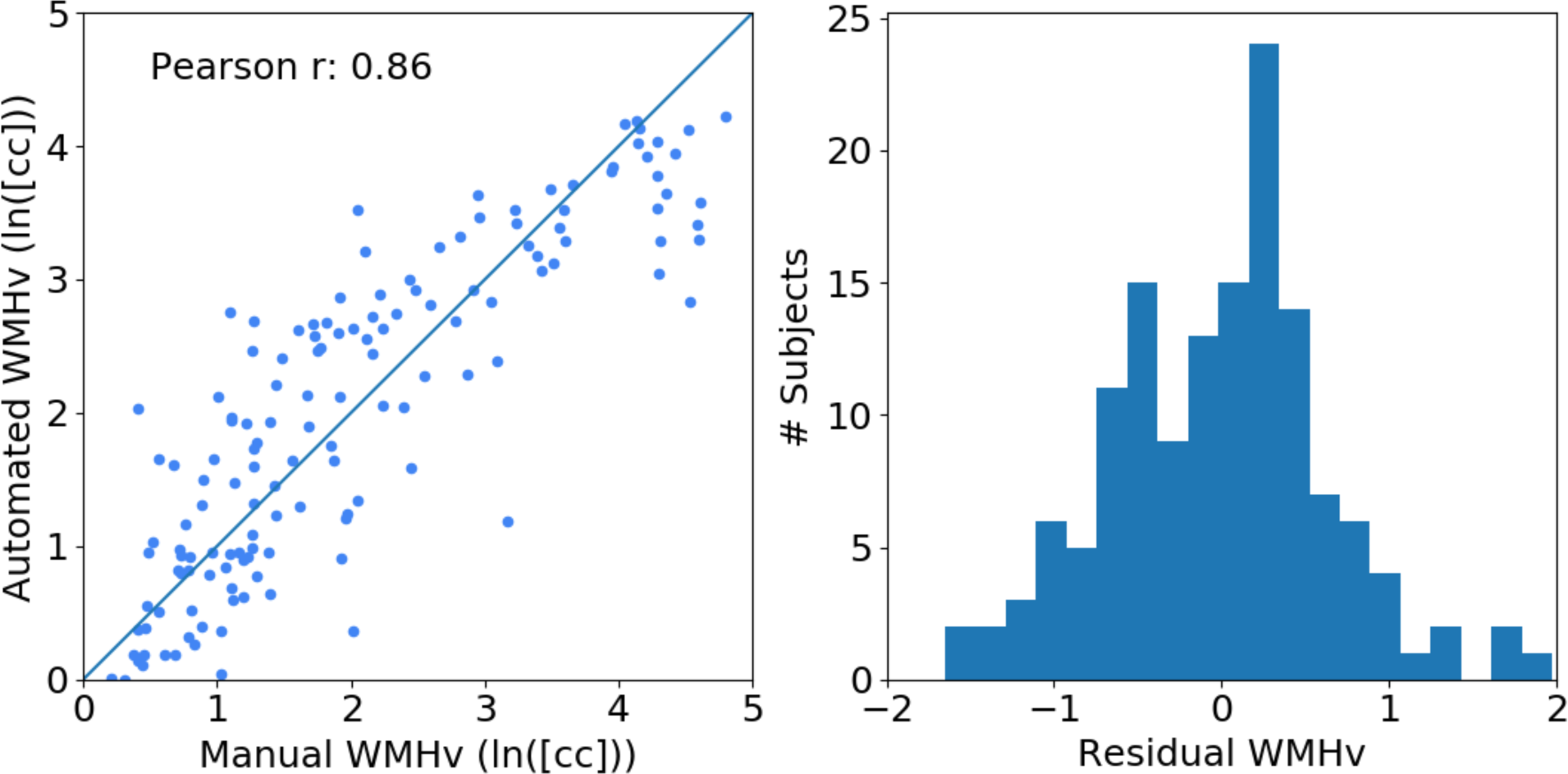
Evaluation of automated and manual WMHv (natural log-transformed). Left: Scatter-plot between automatically and manually determined WMHv (Pearson r = 0.86). The solid line indicates where the automated volume equals the manual volume. Right: Histogram of residuals.

When assessing the residuals of the volume comparison between manual and automated volumes with respect to biases due to differences in age and sex, we did not find any trends in the data. These results suggest that the algorithm can accurately segment WMH in clinical data, facilitating WMH analysis in the entire MRI-GENIE data set.

### Quality control in the MRI-GENIE image set

In total 107 scans (4% of the data) are identified as potential outliers based on assessment of the modified z-score (in- and through-plane resolutions) and those with low slice numbers. Visual inspection identifies 10 incorrectly flagged scans. The remaining 97 scans had a low number of slices (≤3), were different sequences (such as MR angiography, DWI or T1), or included large motion artifacts. These 97 scans (3% of the data) were excluded, leaving 2684 scans for the remaining analysis.

The QC of the automatic analysis pipeline identified potentially erroneous segmentation based on brain extraction. We identified 32 (1%) scans as potential outliers, based on the modified z-score assessment for each site. Further investigation revealed that 3 scans (9% of outliers) were considered outliers due to large motion artefacts, which resulted in removal of large areas of the brain during brain extraction. The majority (n=26; 81% of outliers) were images with a different slice direction (25 coronal scans and 1 sagittal scan). Other problems included wrong acquisition contrast or small brain extraction errors, where parts of the skull and/or eyes were included in the brain mask.

Model (1) of the association of brain volume with age yielded an estimated slope of −2.4 cc/year and an offset of 1,630.8 cc using linear regression. The standard deviation of the residual was 167.5 cc. Figure 8 reports the extracted brain volumes for all subjects, the association of brain volume with age, including two standard deviation for outlier detection. The assessment of how many subjects fall outside the two standard deviations resulted in 163 (6%) flagged subjects.

**Figure 8:**
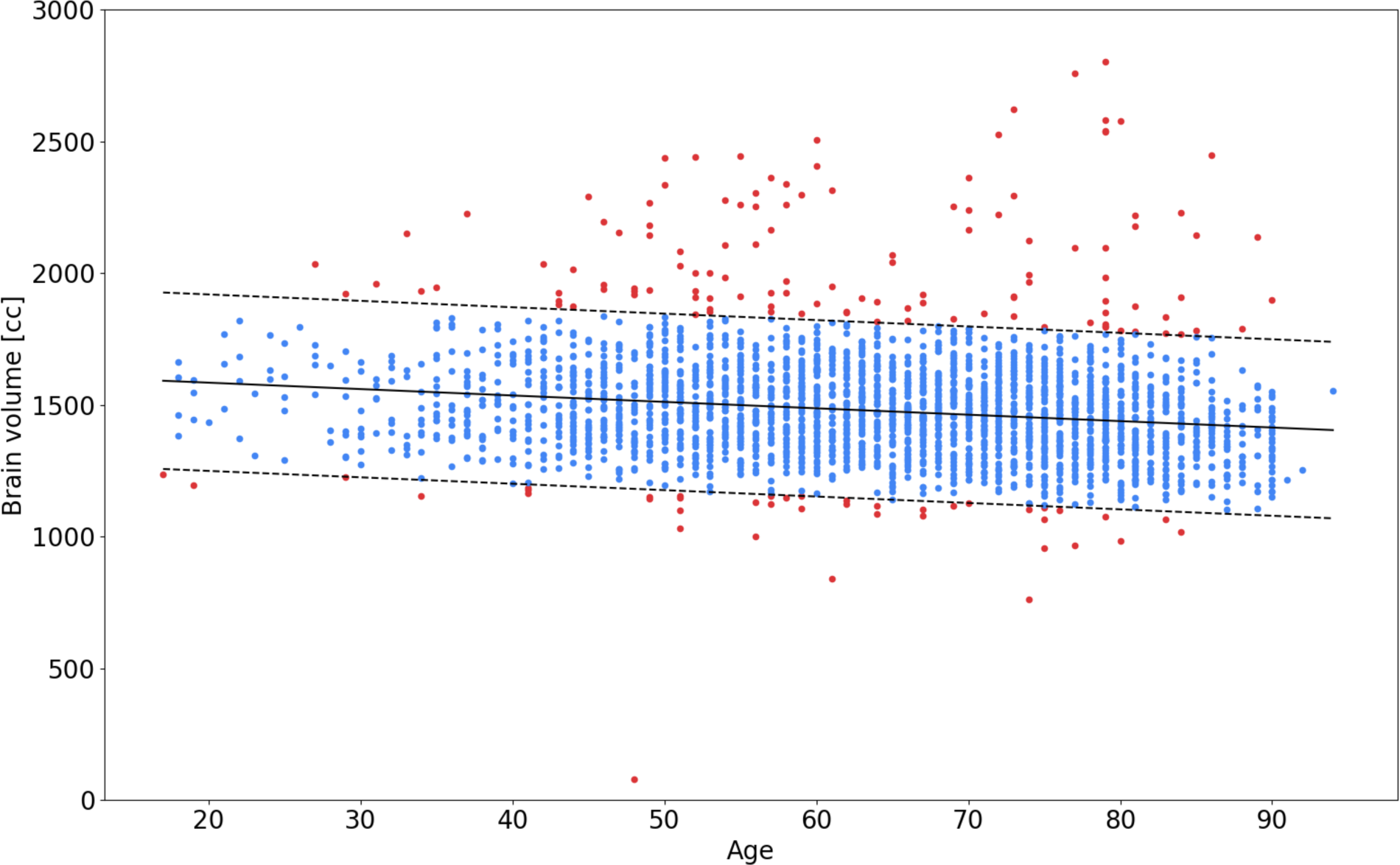
Association of brain volume with age, estimated using automated brain extraction via Neuron-BE for each subject. The solid black line is the estimated linear trend in brain volume with age. Dashed lines represent 2 standard deviation differences from the linear trend, used for outlier (red) detection.

All previously identified outliers in the per-site analysis were also included in this set. Manual assessment of the flagged images showed that 8 (5%) of these scans present with large motion artefacts and 62 (38%) of scans showed small errors, such as incomplete brain extraction. Of all flagged subjects, 10 (6%) were unnecessarily labelled as outliers. The remaining 83 (51%) of scans have wrong acquisition direction (axial requested, sagittal (3; 2%) and coronal (72; 44%) provided, resulting in insufficient stripping of the neck) and other issues (8; 5%). Figure 9 reports site-specific distributions of the estimated brain volume for each imaging site in the study and indicates the flagged images using the site-based and age-based outlier detection.

**Figure 9:**
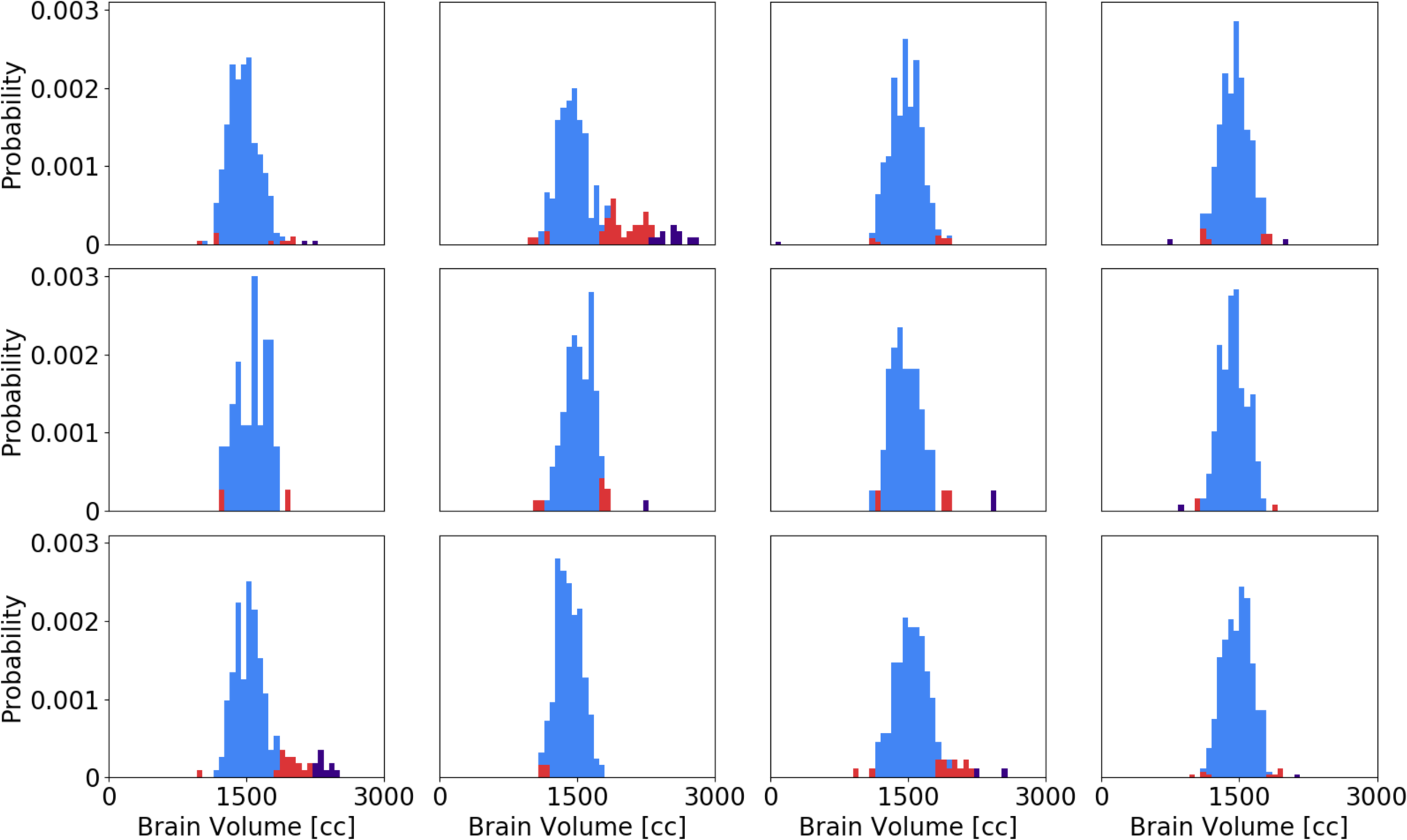
Site-specific distribution of the estimated total brain volume. Red and purple indicate outliers detected using a site-specific and age-based outlier detection, respectively. All site-specific outliers were also identified by the age-based method.

Overall, 250 subjects (9% of the data) were deemed to be outliers and removed from the analysis. The remaining 2,533 subjects (91% of the data) were used to extract WMHv for analysis. As the last QC step, three experts selected two subjects per site at random to sample and manually assess the quality automated WMH outlines. Visual assessment by the raters suggests good agreement of what they would expect to be outlined in these scans.

### WMHv analysis in MRI-GENIE image set

Figure 10 shows the distribution of WMH for each site, in comparison to the global WMH distribution. All sites follow similar trends compared to the pooled WMHv distribution.

**Figure 10:**
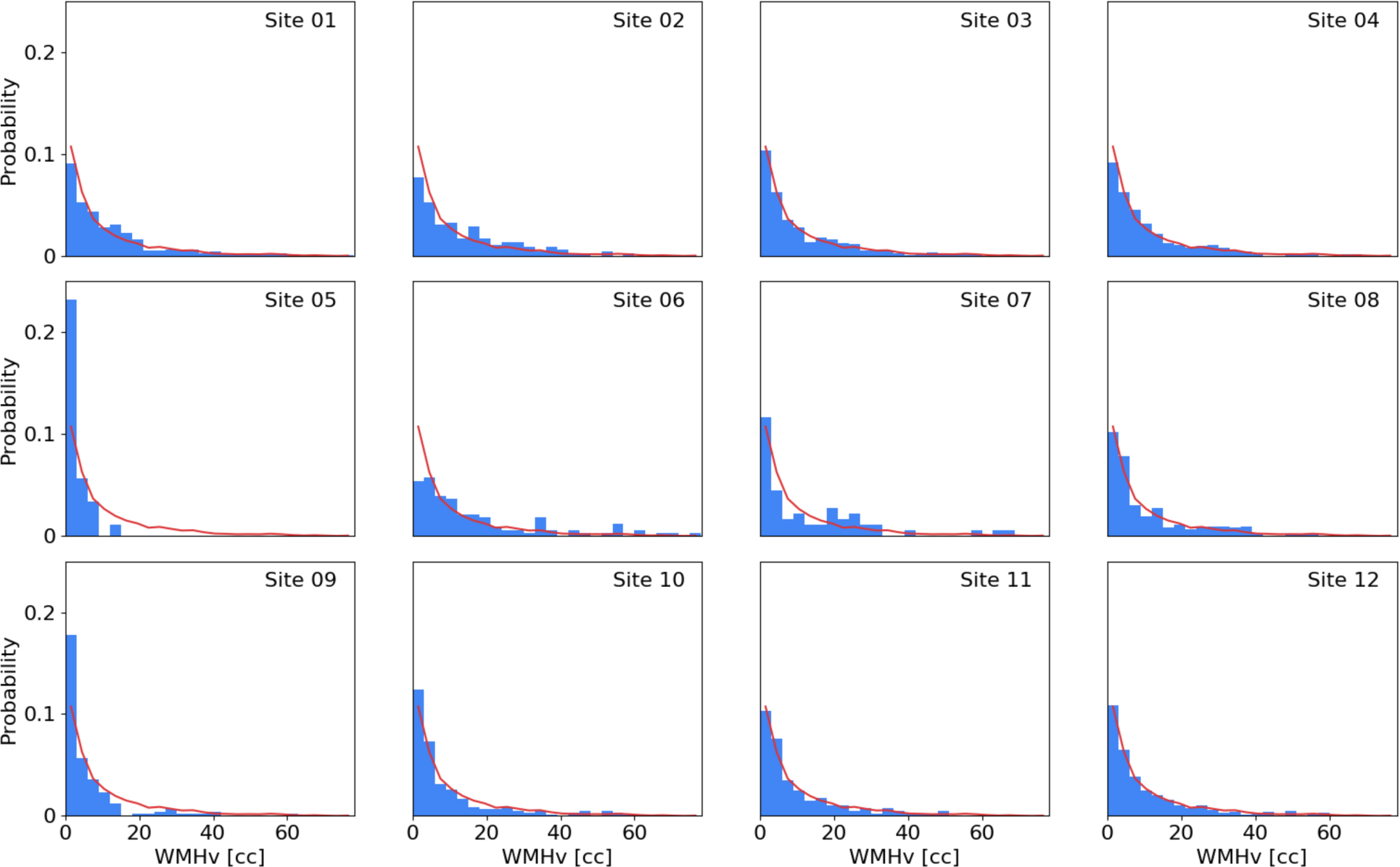
WMHv distributions per individual MRI-GENIE sites (blue histogram), as well as distribution of the combined 12 sites (red line).

Figure 11 shows the age-dependent association of the WMHv in the MRI-GENIE cohort, as well as each individual site. Linear regression results in an estimated slope of 0.051 ln(cc)/year, corresponding to an increase of 0.950 cc/year. We find higher deviation from the pooled estimate for sites with less subjects. *Table 1* summarizes the results stratified by site.

**Table 1:** Summary of quantitative results in this study with the cohort as a whole and stratified by site. N is the number of subjects remaining after QC in relation to initial number of subjects (N_total_). The corresponding characteristics (mean ± standard deviation) include age, brain volume, WMHv and association of WMHv with age, characterized by parameter m in equation (1).

| Site | N/$N_{total}$ | Age (years) | Brain volume (cc) | ln(WMHv (cc)) | Association ln(WMHv (cc)) ~ age (years) |
| --- | --- | --- | --- | --- | --- |
| **All** | 2533 | 63.38±14.58 | 1471.65±150.86 | 1.66±1.35 | 0.051±0.001 |
| **01** | 335/351 | 64.85±15.00 | 1465.38±139.58 | 1.84±1.27 | 0.047±0.002 |
| **02** | 150/202 | 65.21±13.49 | 1444.81±144.42 | 2.06±1.25 | 0.051±0.002 |
| **03** | 448/452 | 65.06±14.36 | 1471.24±151.98 | 1.70±1.30 | 0.051±0.001 |
| **04** | 241/253 | 61.78±13.82 | 1451.94±144.94 | 1.85±1.12 | 0.038±0.001 |
| **05** | 59/61 | 42.59±5.80 | 1451.94±144.94 | 0.45±1.09 | 0.065±0.007 |
| **06** | 110/120 | 70.15±11.10 | 1512.19±139.58 | 2.30±1.13 | 0.042±0.003 |
| **07** | 60/64 | 64.60±15.71 | 1463.68±153.96 | 1.56±1.91 | 0.081±0.002 |
| **08** | 208/289 | 63.89±12.91 | 1439.15±140.96 | 1.69±1.33 | 0.049±0.002 |
| **09** | 159/188 | 52.28±11.68 | 1514.68±142.48 | 0.81±1.54 | 0.071±0.002 |
| **10** | 202/210 | 60.43±13.63 | 1420.39±131.28 | 1.55±1.24 | 0.052±0.002 |
| **11** | 127/148 | 62.71±13.01 | 1523.09±157.05 | 1.62±1.31 | 0.047±0.003 |
| **12** | 434/445 | 67.12±14.45 | 1483.62±152.30 | 1.62±1.33 | 0.048±0.001 |

**Figure 11:**
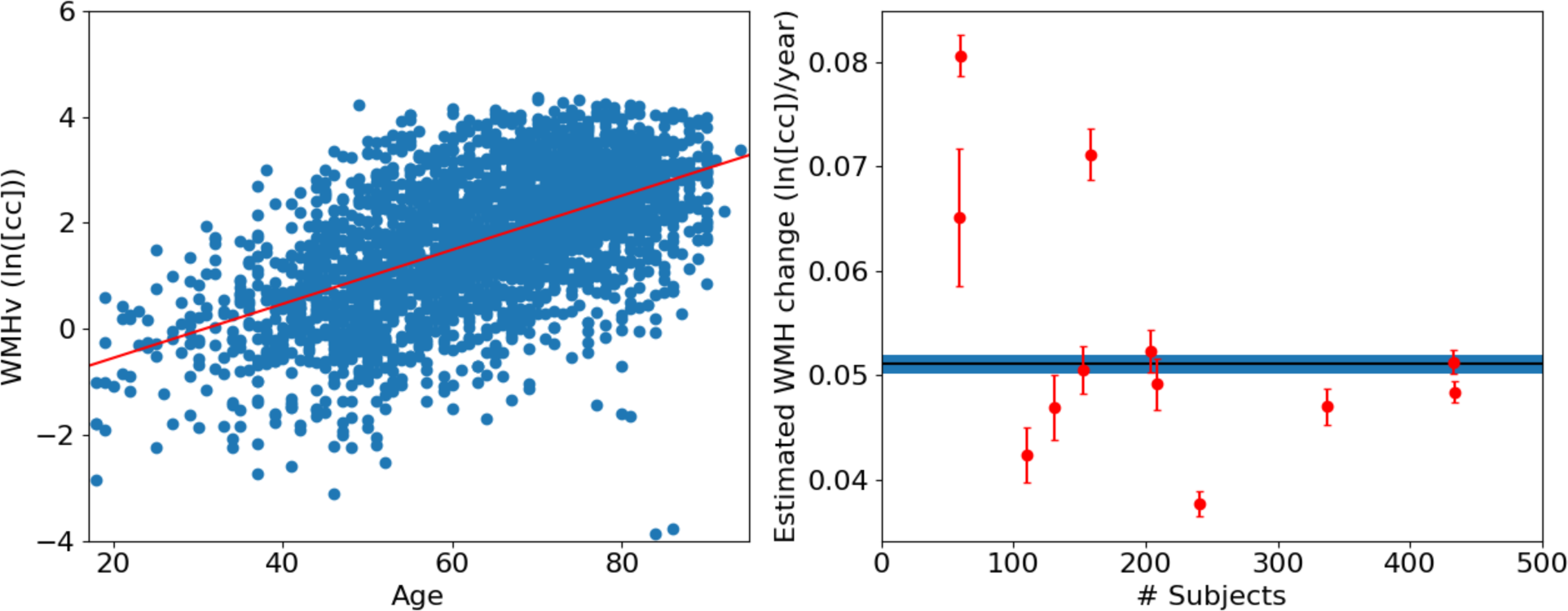
Association of WMHv with age. Left: Regression of log-transformed pooled WMHv from all sites. Right: Association of WMHv against the number of subjects per site. Error bars for each site are computed using a 10-fold split of the data for each site and using a leave-one-fold-out approach to estimate the standard deviations of the coefficient of change. The solid blue bar represents the estimate and standard deviation using all subjects.

## Discussion

Automatic WMH segmentation and WMHv quantification in clinical populations is crucial for elucidating mechanisms, biomarkers and genetic effects in complex diseases such as stroke. In this work, we developed, validated, and implemented a fully automatic pipeline for segmenting WMH in clinical FLAIR images of AIS patients admitted to the hospital. We applied the pipeline to a large-scale international multi-site clinical dataset and investigated associations of WMHv with age.

We assessed the efficacy of the presented pipeline in a validation set of 144 images. WMHv computed from automatic segmentations shows good agreement with that estimated from manual segmentations (ICC=0.84; Pearson *r=0.86*). We find a larger disagreement in scans with higher WMH burden. Visual assessment revealed that the automatic WMH segmentation appears to undersegment in these cases. A likely reason is an underrepresentation of subjects with high WMH burden in the training set of the algorithm. In the 2,533 subjects that passed QC, fewer than 9% of WMHv were estimated to be above 20cc. Future work may include augmenting training of the automatic WMH segmentation by including more of the high-burden WMH cases.

A key contribution of this paper is the preprocessing steps of the analysis pipeline and QC of clinical images. We presented a deep-learning based brain extraction algorithm, which outperforms two of the commonly used and publicly available methods. Outliers due to extreme motion/ghosting artefacts can be identified in the results of our method, enabling this as a QC step of preprocessing and the overall scan quality in large clinical cohorts. We presented two methods of using the estimated brain volumes for QC, where the estimation of age-dependent total brain volume in our cohort demonstrated to be a more rigorous QC criterion compared to site-based modified z-score estimates. We flagged scans as potential erroneous results in the preprocessing, and performed manual assessment. Future improvements in brain extraction could include allowing for both sagittal and coronal scans as inputs, as well as accounting for motion artefacts. However, as WMH burden is clinically best appreciated on axial slices, this might not be beneficial for assessment of WMHv.

We used an intensity normalization step, which is particularly important when working with data acquired across various sites and time, and necessary for intra-site analyses. We showed its efficacy on our validation set where average intensities of white matter were successfully normalized to the same value across subjects and sites.

Applying our pipeline to a large clinical multi-site cohort, we extracted WMHv automatically from 2,533 subjects. Random sampling and assessment of WMH outlines showed good results when assessed by expert raters. Additionally, the WMHv showed expected, approximately exponential distributions for each site. By assessing changes of WMHv with age in this cross-sectional setting, we observe a general WMHv increase with age. Investigating the estimates on a per-site basis, we see higher deviations from the pooled result in sites with lower number of subjects. In future studies, these sites may benefit from being combined to form bigger cohorts, in order to obtain more appropriate representation of the disease burden.

There are several assumptions of this pipeline that merit further consideration. First, it assumes consistent (axial) imaging across the image set. While WMH is best appreciated on these images, occasionally scans are acquired in the sagittal or coronal directions. Future developments to allow other acquisition directions could help recover these images, although studies employing sagittal or coronal imaging directions have to carefully account for the subsequent uncertainties in WMHv. Second, we employ affine rather than deformable spatial normalization. While our results show good agreement with manual outlines, deformable registration may further improve the efficacy of spatial atlas-based priors. Non-linear registration of clinical images, however, is a difficult problem due to the low image quality (Sridharan et al., 2013), potentially leading to gross registration errors, as well as increased computational cost. Third, the main goal of this study was to automate volumetric analysis of WMHv in hospital cohorts of stroke patients. While we focused on the evaluation of accuracy with respect to volumes using the Pearson correlation coefficient, other study may want to utilize the segmentations directly. There are multiple metrics for evaluating the agreement between automated and manual segmentation (Taha and Hanbury, 2015; Wack et al., 2012), including the outline error rate (OER) or the detection error rate (DER); however no consensus has been reached as to which best describes the performance of algorithm. Importantly, the upper bound as to how good an algorithm performs is given by the inter-rater reliability between expert raters. In this study our rater passed the training with an ICC of 0.92, however, due to high effort intensity of manual outlines, we only had segmentations of one expert rater for comparison and could therefore not estimate this upper bound (outline error rate (OER(median (IQR)): 1.00 (0.78-1.22)); DER(median (IQR)): 0.18 (0.05-0.62)).

The strengths of the presented methodology include: (a) the utilization of a large, multi-site, hospital-based cohort of AIS patients with clinical imaging data, as it is acquired in the emergency room; (b) a novel, brain extraction tool for clinical grade FLAIR images based on Unet architecture; (d) the incorporation of simple, automated QC steps for large scale image processing; (d) the adaptation and evaluation of a WMH segmentation algorithm, also based on Unet architecture, specifically designed for hospital-based imaging of AIS patients; and (e) the estimation and evaluation of the association between WMHv and age in over 2500 AIS patients, including an evaluation based on sample size per imaging site.

In conclusion, the presented WMH segmentation pipeline was demonstrated on highly heterogeneous large-scale multi-site data. We applied it to the MRI-GENIE study to demonstrate that its applicability to real-world clinical data, specifically in the acute stroke phase in the hospital. Our method shows promise for future studies that utilize the vast and often under-utilized clinical data to aid phenotypic studies. The resulting phenotypes are not affected by inter-rater variability, which may help reduce variability in follow-up studies, helping to determine additional risk factors for stroke patients. Large studies of WMHv can, for example, elucidate genetic influences, as well as WMH pattern relating to various diseases and phenotypic variables. This work will enable new avenues of research and help advance current knowledge of risks and outcomes in AIS patients.

## Acknowledgements

This project has received funding from the European Union’s Horizon 2020 research and innovation programme under the Marie Sklodowska-Curie grant agreement No 753896 (M.D. Schirmer). This study was supported by NIH-NINDS (MRI-GENIE: R01NS086905 – PI N. Rost, K23NS064052, R01NS082285 – N. Rost, SiGN: U01 NS069208 – J. Rosand, S. Kittner, R01NS059775, R01NS063925, R01NS082285, P50NS051343, R01NS086905, U01 NS069208 – O. Wu), NIH NIBIB (P41EB015902 – P. Golland).

---

**Table A.1:** MRI-GENIE patient cohort. Statistically significant group differences between sites were assessed using ANOVA (age) and χ^2^ tests (sex and prior stroke). All tests between the individual sites were found to be significant (p<0.001). For the validation set only prior stroke was found to be statistically significant (p<0.01), when compared to the remainder of all subjects.

| Site | N | Mean (sd) age | Sex (% male) | % prior stroke |
| --- | --- | --- | --- | --- |
| All | 2783 | 63.28 (14.70) | 61.0 | 10.6 |
| 01 | 351 | 65.32 (15.07) | 61.3 | 0 |
| 02 | 202 | 64.59 (14.44) | 48.5 | 28.7 |
| 03 | 452 | 64.89 (14.48) | 64.4 | 14.5 |
| 04 | 253 | 61.98 (13.83) | 62.1 | 12.6 |
| 05 | 61 | 42.08 (6.59) | 78.7 | 0 |
| 06 | 120 | 70.05 (10.93) | 62.5 | 17.6 |
| 07 | 64 | 64.23 (16.12) | 53.1 | 25 |
| 08 | 289 | 63.58 (13.41) | 72.0 | 19.7 |
| 09 | 188 | 52.38 (11.58) | 60.1 | 9.6 |
| 10 | 210 | 60.54 (13.85) | 52.4 | 9.5 |
| 11 | 148 | 62.71 (13.09) | 60.8 | 0 |
| 12 | 445 | 67.02 (14.63) | 58.2 | 1.8 |
| Validation set | 144 | 62.01 (16.85) | 59.7 | 18.2 |

## Footnotes

1 Neuron-BE is implemented by building on the open-source neuron library found at http://github.com/adalca/neuron (Dalca et al., 2018)

